# A Framework to Incorporate D-trace Loss into Compositional Data Analysis

**DOI:** 10.1101/464982

**Authors:** Shun He, Minghua Deng

## Abstract

The development of high-throughput sequencing technologies for 16S rRNA gene profiling provides higher quality compositional data for microbe communities. Inferring the direct interaction network under a specific condition and understanding how the network structure changes between two different environmental or genetic conditions are two important topics in biological studies. However, the compositional nature and high dimensionality of the data are challenging in the context of network and differential network recovery. To address this problem in the present paper, we proposed a framework to incorporate the data transformations developed for compositional data analysis into D-trace loss for network and differential network estimation, respectively. The sparse matrix estimators are defined as the minimizer of the corresponding lasso penalized loss. This framework is characterized by its straightforward application based on the ADMM algorithm for numerical solution. Simulations show that the proposed method outperforms other state-of-the-art methods in network and differential network inference under different scenarios. Finally, as an illustration, our method is applied to a mouse skin microbiome data.

**Author summary:** Inferring the direct interactions among microbes and how these interactions change under different conditions are important to understand community-wide dynamics. The compositional nature and high dimensionality are two distinctive features of microbial data, which invalidate traditional correlation analysis and challenge interaction network estimation. In this study, we set up a framework that combines data transformation with D-trace loss to infer the direct interaction network and differential network from compositional data. Simulations and real data analysis show that our proposed methods lead to results with higher accuracy and stability.

## 1 Introduction

Microbes play critical roles in Earth’s biogeochemical cycles [8] and impact the health of humans significantly [24]. Understanding interactions among microbes under a specific condition is a key research topic in microbial ecology [17]. Bandyopadhyay *et al.* [3] also showed that these interactions can change under various environmental or genetic conditions. With the development of high-throughout sequencing technology, 16s rRNA gene sequences can be amplified, sequenced, and grouped into common Operational Taxonomic Units (OTUs), and as a result, microbial abundance information can be obtained for further exploration [27]. One of the major challenges is to discover associations among microbes and how these associations change under different conditions, which could in turn help us to unravel the underlying interaction network and offer an insight into community-wide dynamics.

Correlation analysis is commonly used to infer the interaction network for absolute abundance data. However, applying traditional correlation analysis to compositional data, as only representative of relative abundances of microbial species, may yield spurious results [1, 14]. Recent methods, such as SparCC [14], CCREPE [11, 12], REBACCA [2] and CCLasso [9], have been proposed to address compositional bias and infer the correlation network of microbe communities. However, pairwise correlations contain both direct and indirect interactions, and correlations may arise when microbes are connected indirectly [18]. Thus, the conditional dependence network describing direct interactions is often more intrinsic and fundamental [10, 15].

For absolute abundance, conditional dependence networks are frequently modeled as Gaussian graphical models where direct interactions are correspond to the support of precision matrix [19, 26]. Meinshausen and Bühlmann [20] proposed a neighborhood selection approach to recover the precision matrix row-by-row by fitting a lasso penalized least square regression model [25]. Yuan and Lin [29] derived the likelihood for Gaussian graphical models and suggested using the maxdet algorithm to compute the corresponding lasso penalized estimator. Friedman *et al.* [13] developed a more efficient algorithm called the graphical lasso. Zhang and Zou [30] proposed a new loss function called D-trace loss and introduced a sparse precision matrix estimator as the minimizer of lasso penalized D-trace loss. Several methods have been proposed to infer the direct interaction network from compositional data. Biswas *et al.* [5] suggested learning the direct interactions from compositional data with a Poisson-multivariate normal hierarchical model called MInt. Kurtz*et al.* [18] proposed a method called SPIEC-EASI, which combines centered log-ratio (clr) transformation [1] for compositional data with the neighborhood selection approach [20] or graphical lasso [13] to estimate the precision matrix. Similar to the idea of Yuan and Lin [29], Fang *et al.* [10] first derived likelihood with compositional data for Gaussian graphical models and then estimated the precision matrix with a lasso penalized maximum likelihood method called gCoda. In this paper, we proposed a framework to incorporate clr transformation [1] into D-trace loss [30] to estimate the precision matrix from compositional data.

Biological networks often vary according to different environmental or genetic conditions [3]. Understanding how networks change and estimating differential networks are important tasks in biological studies. In recent years, researchers have actively sought methods of estimating differential networks for absolute abundance data. Chiquet *et al.* [6], Guo *et al.* [16] and Danaher *et al.* [7] estimated the precision matrices and their differences jointly by penalizing the joint log-likelihood with different penalties. Zhao *et al.* [31] developed a .*ℓ*_1_-minimization method for direct estimation of differential networks, which does not require sparsity of precision matrices or their separate estimation. Yuan *et al.* [28] proposed a new loss function called DTL based on D-trace loss [30] to estimate the precision matrix difference directly. In this paper, we also extended our framework to incorporate clr transformation [1] into DTL [28] to estimate the differential network from compositional data.

The remainder of the paper is organized as follows. In Section 2, we introduce our framework to incorporate clr transformations for compositional data analysis into D-trace loss, thereby enabling us to estimate both direct interaction network and differential direct interaction networks from compositional data, respectively. In Section 3, the performance of our method was evaluated and compared with other state-of-the-art methods under various simulation scenarios. In Section 4, the proposed methods are illustrated with an application to a mouse skin microbiome data.

## 2 Materials and methods

### 2.1 Compositional Data and clr Transformation

We begin with some notations and definitions for convenience. For a *p* × *p* matrix *X* = (*X_ij_*) ∈ 𝓡^*p*×*p*^, its transposition, trace and determinant are denoted as *X*^T^, *tr*(*X*) and det *X*, respectively. Let
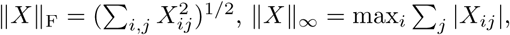 |*X*|_1_ = max_*j*_ ∑_*i*_ |*X_ij_*|, ∥*X*∥_1_ = ∑_*i*,*j*_ |*X_ij_*|, and ∥*X*∥_1,off_ = ∑_*i*≠*j*_ |*X_ij_*| be the Frobenius norm, ∞-norm, 1-norm, *ℓ*_1_-norm and off-diagonal *ℓ*_1_-norm of *X*. Denote by *vec*(*X*) the *p*^2^-vector from stacking the columns of *X*, and *X* ≻ 0 means that *X* is positive definite. For two matrices *X*, *Y* ∈ 𝓡^*p*×*p*^, let *X* ⊗ *Y* be the Kronecker product of *X* and *Y* . We use 〈*X*, *Y*〉 to denote *tr*(*XY*^T^) throughout this paper.

Suppose that there are *p* microbe species and that their absolute abundances are
***z*** = (*z*_1_, *z*_2_, …, *z_p_*) respectively. However, instead of absolute abundances, it is often the case that only the relative abundances (or compositions) ***x*** = (*x*_1_, *x*_2_, …, *x_p_*), where

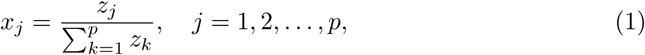

can be observed in real experiments. If the log-transformed absolute abundances ln ***z*** follow a multivariate Gaussian distribution with mean ***μ*** and nonsingular covariance matrix Σ, the precision matrix Θ = Σ^−1^ depicts the direct interaction network among microbial specials since ln *z_i_* and ln *z_j_* are conditionally independent given other components of ***z*** if and only if Θ*_ij_* = 0 [15]. Moreover, we can describe this direct interaction network with an undirected graph if we represent the *p* microbe species with *p* vertices and connect the conditionally dependent species pairs.

Log-ratios
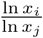
[1] are commonly used in compositional data analysis, since the simple equality
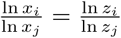 serves as the bridge between the compositions and the unobserved absolute abundances [18]. Aitchison [1] also proposed a statistically equivalent centered log-ratio (clr) transformation. The clr transformation matrix is
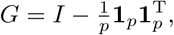 where **1**_*p*_ is a *p*-dimensional all-ones vector and *I* is identity matrix. Applying the clr transformation and using ln ***x*** = ln ***z*** − **1**_*p*_ ln *s* and *G***1**_*p*_ = **0**_*p*_, it follows that

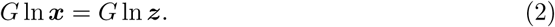

Denoted by Σ_ln ***x***_ the covariance matrix of the log-transformed relative abundances, we have

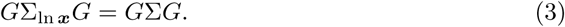

Similarly, equation (2) and (3) establish a bridge between the observed relative abundances and the unobserved absolute abundances. SPIEC-EASI [18] assumes that *G*Σ_ln ***x***_ *G* serves as a good approximation of Σ since *G* − *I* ≈ 0 when *p* ≫ 0, and apply the neighborhood selection approach [20] or graphical lasso [13] to the clr-transformed relative abundances for precision matrix estimation.

### 2.2 CDTr : Compositional Network Analysis with D-trace Loss

From the empirical loss minimization perspective, SPIEC-EASI is not the most natural and concise because of the approximation and the log-determinant term in graphical lasso [13]. In this section, we introduce an innovative framework to estimate the direct interaction network from compositional data with D-trace loss. The new D-trace loss for compositional data (CDTr loss) is proposed as

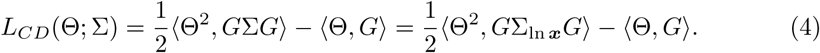

We can view the CDTr loss as an analogue of the D-trace [30] loss
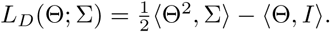 The meaning of incorporating clr transformation into the original D-trace loss is to avoid the unobserved absolute abundance and account for the compositionality. If we know the absolute abundance data, we can simply substitute the finite sample estimator of Σ sample estimator of (denoted by
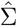) into D-trace loss and estimate the precision matrix Θ with the corresponding lasso penalized estimator. However, for relative abundances or compositional data, only the finite sample estimator of Σ_ln ***x***_ (denoted by
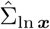) is available, instead of the finite sample estimator of Σ. Thanks to the clr transformation and the bridge equation (3), we can estimate *G*∑*G* with
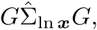 even though
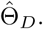 is not available.

It is easy to check that CDTr loss can be written as

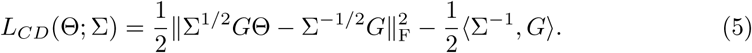

To ensure that Σ^−1^ minimizes *L_CD_*, namely Σ^1/2^*G*Θ − Σ^−1/2^*G* = 0 when Θ = Σ^−1^, we need the following exchangeable condition:

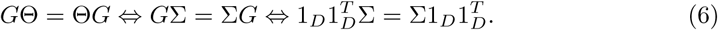

Denote by *σ_ij_* and *ρ_ij_* the covariance and correlation between ln *z_i_* and ln *z_j_*, respectively. Then, the exchangeable condition is equivalent to ∑*_l_ σ_il_* = ∑*_l_ σ_jl_* for all *i*, *j* = 1, 2, …, *p*, which is similar to the assumption ∑_*l*≠*i*_ *σ_il_* = 0, *i* = 1, 2, …, *p* in SparCC [14]. If the variances *σ_ii_*, *i* = 1, 2, …, *p* are all the same, then the exchangeable condition simplifies to ∑_*l*≠*i*_ *ρ_il_*, *i* = 1, 2, …, *p* are all the same, which implies that the average correlation with other species is nearly the same for each specie. Analogously, the assumption in SparCC simplifies to ∑_*l*≠*i*_ *ρ_il_* = 0, *i* = 1, 2, …, *p*, which implies that the average correlations are very small. In the numerical experiments of section 3, we show that CDTr still performs well, even when the exchangeable condition does not hold.

In practical applications, we use the empirical version of CDTr loss as

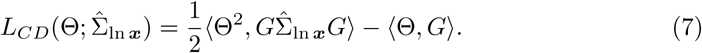

Since most species do not interact directly when the number of species *p* is large, we further assume that the direct interaction network, or Θ, is sparse, which also helps to solve the under-determined problem caused by compositionality and dimensionality [9, 10, 29]. We employ the commonly used *ℓ*_1_ penalty [25, 29, 30] to handle the sparse assumption, and our sparse estimator of the precision matrix Θ is proposed as

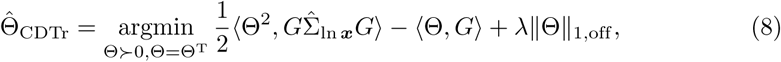

where λ ≥ 0 is the tuning parameter for the tradeoff between the model fitting and the sparsity of
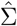 Following the idea of Zhao *et al.* [31], the tuning parameter is selected by minimizing the Bayesian Information Criterion (BIC) [22] as

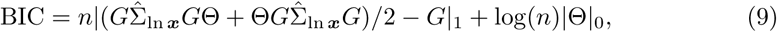

where |Θ|_0_ is the number of non-zero elements in the upper-triangle of Θ, and *n* is the sample size.

Zhang and Zou [30] developed an efficient algorithm based on alternating direction methods [21] for the solution of penalized D-trace loss estimator. We can simply replace
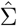 and *I* in D-trace loss with
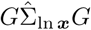 and *G* in our CDTr loss and use the algorithm of Zhang and Zhou [30] for the numerical solution of (8). Following the idea of Zhang and Zhou [30] and Scheinberg *et al.* [21], we introduce two new matrices, Θ_0_ and Θ_1_. The augmented Lagrangian function of (8) are considered, and Λ_0_, Λ_1_, *ρ* are Lagrangian multipliers. The steps of the ADMM algorithm for the lasso penalized CDTr loss estimator are summaried in Algorithm 1. The definitions of matrix operators

#### Algorithm 1

The algorithm for lasso penalized CDTr loss estimator.

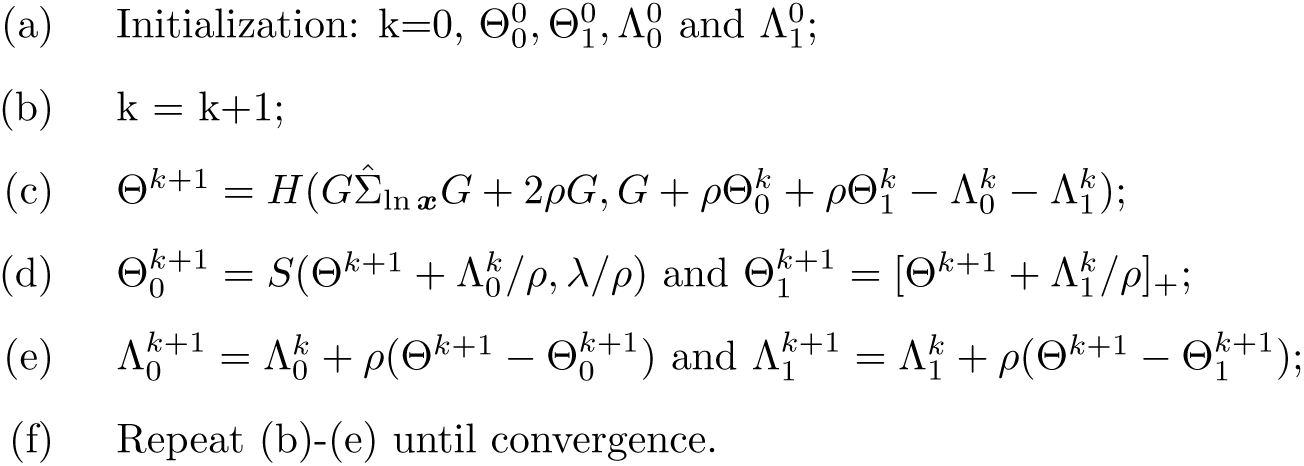

*H*(*X*), *S*(*X*) and [*X*]_+_ are listed in S1 Appendix.

### 2.3 DCDTr : Differential Compositional Network Analysis with D-trace Loss

Consider that the absolute abundances of *p* microbe species become
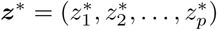 under another condition and that the relative abundances are
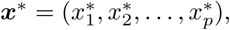 respectively. Similarly, we assume ln ***z*^∗^** ∼ 𝓝 (***μ*^∗^**, Σ^∗^). Thus, we want to estimate the difference between direct interaction networks under different conditions, i.e., the resultant differential network Δ = Σ^∗−1^ − Σ^−1^.

A straightforward approach to estimate Δ is to estimate Σ^−1^ and Σ^∗−1^ separately and then subtract the estimates under the key assumption that both precision matrices are sparse. However, a more reasonable assumption is that the difference between the precision matrices are sparse, not that both matrices are sparse, since direct interactions may not be sparse while the changes under different conditions are often sparse [28]. Therefore, we proposed a new loss function for differential network estimation with compositional data (DCDTr loss) to estimate Δ directly, under the assumption that the differential network Δ is sparse. The DCDTr loss is proposed as

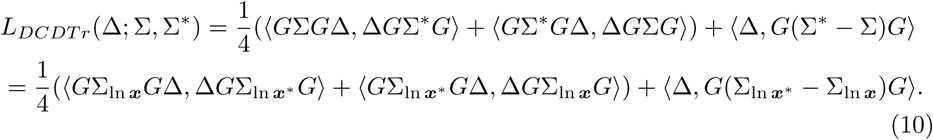

Similarly, our DCDTr loss can be regarded as an analogue to the DTL loss
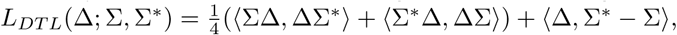 which is proposed by Yuan *et al.* [28] to estimate the differential network Δ when the absolute abundances are known. Again, our DCDTr loss takes the advantage of the bridge equation (3) to avoid the unobserved absolute abundance and account for the compositionality. From another perspective, we can arrive at our DCDTr loss (10) by substituting the approximation Σ ≈ *G*Σ_ln **x**_ *G*, Σ^∗^ ≈ *G* Σ_ln ***x***^∗^_ *G* into DTL loss. In the numerical experiments of section 3, we also investigated the performance of procedures which combine the approximation Σ ≈ *G* Σ_ln ***x***_ *G*, Σ^∗^ ≈ *G*Σ_ln ***x***^∗^_ *G* with other methods for differential network estimation, including the *ℓ*_1_-minimization method [31] for direct estimation of differential networks and joint graphical lasso (FGL, GGL) [7] for joint estimation of precision matrices. The detailed formulas are left in **??**.

Under the exchangeable condition *G*Σ = Σ*G* and *G* Σ^∗^ = Σ^∗^ *G*, it is easy to check that

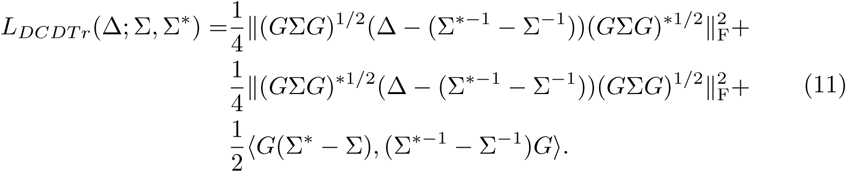

Obviously, Δ = Σ^∗−1^ − Σ^−1^ is a minimizer of our DCDTr loss *L_DCDTr_* . In practical applications, we incorporate the finite sample estimators of Σ, Σ^∗^ and *ℓ*_1_ penalty into DCDTr loss, and our sparse estimator for the differential network Δ is proposed as

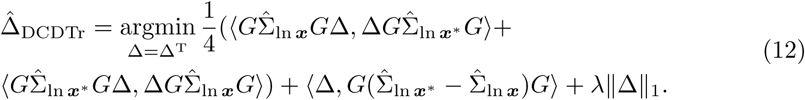

The tuning parameter λ is selected by minimizing the Bayesian Information Criterion (BIC) [22, 28, 31] as

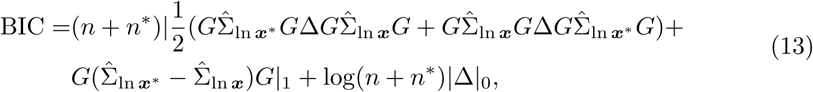

where |Δ|_0_ is the number of non-zero elements in the upper-triangle of Δ, and *n* and *n*^∗^ are the sample size.

Taking advantage of the algorithm developed by Yuan *et al.* [28] for the numerical solution of lasso penalized DTL loss estimator, the algorithm for the numerical solution of (12) is straightforward, essentially because we can simply replace
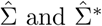
in DTL loss with
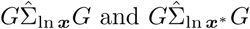 in our DCDTr loss. Following the idea of Yuan *et al.* [28], we introduce three new matrices Δ_1,2,3_ and Lagrangian multipliers Λ_1,2,3_, *ρ* for the solution of (12). The steps of the ADMM algorithm for the lasso penalized DCDTr loss estimator are presented in Algorithm 2. The definitions of matrix operators *K*(*X*) and *S*(*X*) are listed in S1 Appendix.

## 3 Numerical Results

In this section, we conduct several numerical experiments under different settings and compare them with other state-of-the-art methods. Given mean ***μ**_p_* and precision matrix Θ, we first generate the log-transformed absolute abundance ln ***z**_i_* = (ln *z*_*i*1_, ln *z*_*i*2_, …, ln *z_ip_*) with the multivariate normal distribution 𝓝_*p*_(***μ**_p_*, Θ^−1^), and then the relative abundances are
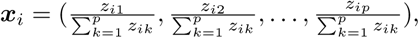
*i* = 1, 2, …, *n*. For another given mean
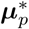 and precision matrix Θ^∗^ under a new condition, the samples
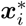, *i* = 1, 2, …, *n* are similarly generated. In the following simulations, we take *p* = 50 and ***μ**_p_* sampled from the uniform distribution 𝓤_*p*_(−0.5, 0.5).

### Algorithm 2

The algorithm for lasso penalized DCDTr loss estimator.

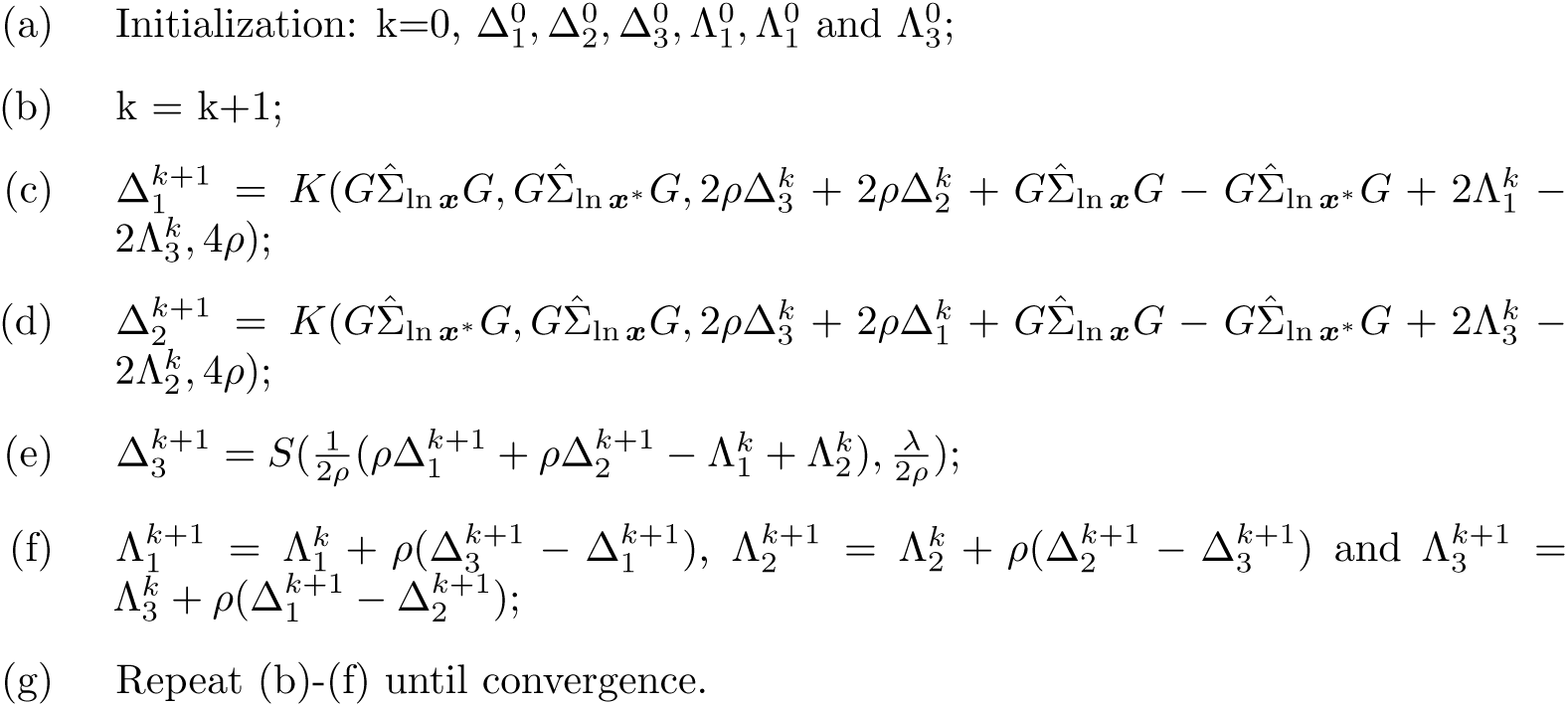

### 3.1 Simulations for CDTr Loss

To investigate the performance of CDTr loss and the influence of the exchangeable condition, we considered the following network structures for Θ.

#### 1. Band graph

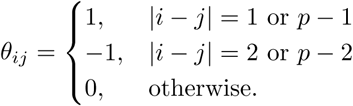

#### 2. Cluster graph

Divide *p* nodes into 5 clusters evenly. The nodes in different clusters are not connected, while the network for each cluster is the same as matrix *C* = (*c_ij_*)_10×10_, where

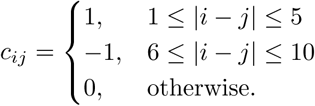

The link strength is uniformly distributed in [*l*, *u*]. To be specific, *θ_ij_* is replaced with *θ_ij_s_ij_*, where *s_ij_* ∼ 𝓤 (*l*, *u*). We take (*l*, *u*) = (0.1, 0.1), (0.05, 0.15) and (0.0.2) separately to study the performance of CDTr loss when the exchangeable condition is satisfied by different degrees. These scenarios are named as Band-exact (Band-e), Band-approx1 (Band-a1), Band-approx2 (Band-a2) and Cluster-exact (Cluster-e), Cluster-approx1 (Cluster-a1), Cluster-approx2 (Cluster-a2), respectively. To obtain a positive definite precision matrix Θ, we first compute the smallest eigenvalue of Θ (denoted by *e*); then the diagonal elements of Θ are set as |*e*| + 0.3. The deviation to the exchangeable condition is measured with *dev* = ∥*G*Σ − Σ*G*∥_F_. The deviations under the aforementioned six scenarios are listed in Table 1. For each combination of the six network structures and four sample sizes *n* = 50, 100, 150, 200, a total of 100 datasets are generated and used to recover the network structure. Three state-of-the-art methods for network recovery are investigated, including gCoda [10], SPIEC(MB) and SPIEC(GL) [18]. We further consider an approximation method called aCDTr, which approximates Σ with *G*Σ_ln ***x**G*_ [18] and employs D-trace loss to estimate Θ = Σ^−1^. Specifically, the estimator of aCDTr is

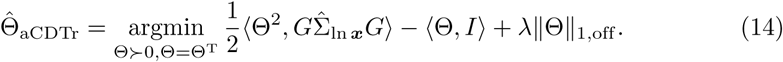

**Table 1.** Deviations from the exchangeable condition under different scenarios.

| Network | Band-e | Band-a1 | Band-a2 | Cluster-e | Cluster-a1 | Cluster-a2 |
| --- | --- | --- | --- | --- | --- | --- |
| $dev$ | 0 | 0.203 | 0.348 | 0 | 0.109 | 0.205 |

The true positive rate and true negative rate are evaluated at different tuning parameters and used to generate the receiver operating characteristic (ROC) curve. We use the area under the curve (AUC) to quantify the ability to recover the true underlying network.

In Table 2, we present the mean AUC scores of the above-mentioned methods under different settings. The mean AUC scores of CDTr and aCDTr are superior to the other three methods in all cases, even when the exchangeable condition does not hold exactly, which implies that CDTr and aCDTr outperform other methods in direct interaction network recovery. Moreover, the mean AUC of CDTr is slightly higher than that of aCDTr, except for the cluster graph and sample size *n* = 50. With increasing deviation, the performance of CDTr and aCDTr decreases, which is reasonable if the exchangeable condition does not exactly hold. Interestingly, the performance for the other three methods also decreases with increasing deviation. For all network structures and methods, the mean AUC scores increase as the sample size increases.

**Table 2.**
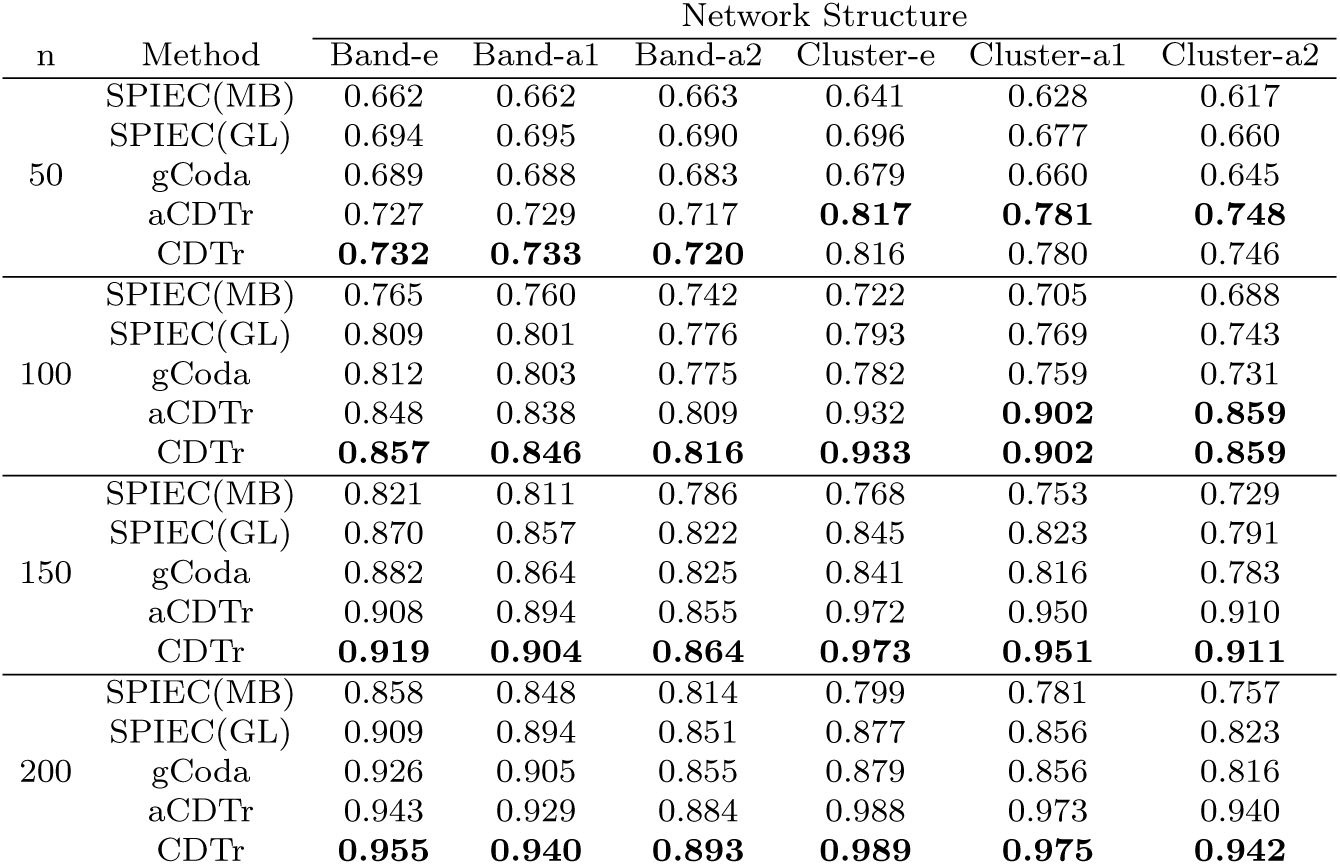
The mean AUC scores of different methods under different settings.

| n | Method | Network Structure |  |  |  |  |  |
| --- | --- | --- | --- | --- | --- | --- | --- |
|  |  | Band-e | Band-a1 | Band-a2 | Cluster-e | Cluster-a1 | Cluster-a2 |
| 50 | SPIEC(MB) | 0.662 | 0.662 | 0.663 | 0.641 | 0.628 | 0.617 |
|  | SPIEC(GL) | 0.694 | 0.695 | 0.690 | 0.696 | 0.677 | 0.660 |
|  | gCoda | 0.689 | 0.688 | 0.683 | 0.679 | 0.660 | 0.645 |
|  | aCDTr | 0.727 | 0.729 | 0.717 | **0.817** | **0.781** | **0.748** |
|  | CDTr | **0.732** | **0.733** | **0.720** | 0.816 | 0.780 | 0.746 |
| 100 | SPIEC(MB) | 0.765 | 0.760 | 0.742 | 0.722 | 0.705 | 0.688 |
|  | SPIEC(GL) | 0.809 | 0.801 | 0.776 | 0.793 | 0.769 | 0.743 |
|  | gCoda | 0.812 | 0.803 | 0.775 | 0.782 | 0.759 | 0.731 |
|  | aCDTr | 0.848 | 0.838 | 0.809 | 0.932 | **0.902** | **0.859** |
|  | CDTr | **0.857** | **0.846** | **0.816** | **0.933** | **0.902** | **0.859** |
| 150 | SPIEC(MB) | 0.821 | 0.811 | 0.786 | 0.768 | 0.753 | 0.729 |
|  | SPIEC(GL) | 0.870 | 0.857 | 0.822 | 0.845 | 0.823 | 0.791 |
|  | gCoda | 0.882 | 0.864 | 0.825 | 0.841 | 0.816 | 0.783 |
|  | aCDTr | 0.908 | 0.894 | 0.855 | 0.972 | 0.950 | 0.910 |
|  | CDTr | **0.919** | **0.904** | **0.864** | **0.973** | **0.951** | **0.911** |
| 200 | SPIEC(MB) | 0.858 | 0.848 | 0.814 | 0.799 | 0.781 | 0.757 |
|  | SPIEC(GL) | 0.909 | 0.894 | 0.851 | 0.877 | 0.856 | 0.823 |
|  | gCoda | 0.926 | 0.905 | 0.855 | 0.879 | 0.856 | 0.816 |
|  | aCDTr | 0.943 | 0.929 | 0.884 | 0.988 | 0.973 | 0.940 |
|  | CDTr | **0.955** | **0.940** | **0.893** | **0.989** | **0.975** | **0.942** |

We further conducted several experiments on the following six representative network structures, without considering the exchangeable condition.

1. *Random graph*: Two nodes are connected with probability 0.1, and the strength is generated from a uniform distribution in [−0.2, − 0.1] ∪ [0.1, 0.2].
2. *Band graph*: Connect pair (*i*, *j*) with strength uniformly distributed in [0.05*m* − 0.3, 0.05*m* − 0.25] ∪ [0.25 − 0.05*m*, 0.3 − 0.05*m*], if |*i* − *j*| = *m*, *m* = 1, 2, 3, 4.
3. *Neighbor graph*: Select *p* points from 𝓤(0, 1) and connect the 5 nearest neighbors for each point with strength sampled from a uniform distribution in [− 0.15, − 0.05] ∪ [0.05, 0.15].
4. *Scale*-*free graph*: A scale-free graph is produced, following the B-A algorithm [4]. The initial graph has two connected nodes, and each new node is connected to only one node in the existing graph with the probability proportional to the degree of the each node in the existing graph. This results in *p* edges in the graph, and the strength between connected nodes is generated from a uniform distribution in [−0.2, − 0.1] ∪ [0.1, 0.2].
5. *Hub graph*: Partition the nodes into 3 disjoint groups evenly and select a node as hub for each group. The hubs are connected with the non-hubs in the same group with strength uniformly distributed in [−0.2, − 0.1] ∪ [0.1, 0.2].
6. *Block graph*: Divide *p* nodes into 5 blocks evenly. Connect pairs in the same block with probability 0.3 and pairs in different blocks with probability 0.1. The strength between connected nodes is uniformly distributed in [−0.2, − 0.1] ∪ [0.2, 0.1].

Similarly, the diagonal elements of Θ are set as |*e*| + 0.3, where *e* is the smallest eigenvalue of Θ. The deviations from the exchangeable condition of these networks are listed in Table 3.

**Table 3.**
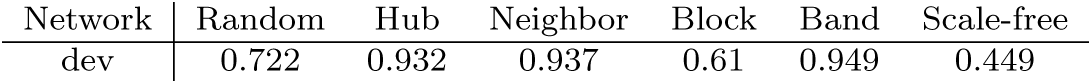
Deviations from the exchangeable condition of six different network structures.

We generated 100 datasets for each setting and used them to estimate the true precision matrix. The mean AUC scores of different methods under different settings are shown in Table 4. We can see that CDTr performs better than other methods in all cases, while the results of aCDTr is comparable to those of gCoda, and the results of SPIEC(MB) and SPIEC(GL) is are worse than the others. Note that we did not consider the exchangeable condition when we set up the networks, implying that CDTr still works, even when the the exchangeable condition does not hold.

**Table 4.** The mean AUC scores of different methods under different settings.

| n | Method | Network Structure |  |  |  |  |  |
| --- | --- | --- | --- | --- | --- | --- | --- |
|  |  | Random | Band | Neighbor | Scale-free | Hub | Block |
| 50 | SPIEC(MB) | 0.630 | 0.615 | 0.599 | 0.671 | 0.647 | 0.613 |
|  | SPIEC(GL) | 0.652 | 0.637 | 0.616 | 0.697 | 0.690 | 0.635 |
|  | gCoda | 0.652 | 0.636 | 0.615 | 0.700 | 0.745 | 0.633 |
|  | aCDTr | 0.677 | 0.664 | 0.636 | 0.728 | 0.748 | 0.660 |
|  | CDTr | **0.681** | **0.670** | **0.641** | **0.729** | **0.757** | **0.664** |
| 100 | SPIEC(MB) | 0.728 | 0.687 | 0.674 | 0.785 | 0.767 | 0.693 |
|  | SPIEC(GL) | 0.758 | 0.712 | 0.697 | 0.809 | 0.812 | 0.723 |
|  | gCoda | 0.766 | 0.717 | 0.703 | 0.811 | 0.866 | 0.729 |
|  | aCDTr | 0.778 | 0.737 | 0.714 | 0.827 | 0.860 | 0.746 |
|  | CDTr | **0.786** | **0.745** | **0.726** | **0.831** | **0.872** | **0.754** |
| 150 | SPIEC(MB) | 0.782 | 0.731 | 0.713 | 0.845 | 0.834 | 0.746 |
|  | SPIEC(GL) | 0.816 | 0.758 | 0.742 | 0.870 | 0.877 | 0.783 |
|  | gCoda | 0.831 | 0.770 | 0.756 | 0.874 | 0.925 | 0.796 |
|  | aCDTr | 0.832 | 0.779 | 0.759 | 0.883 | 0.914 | 0.802 |
|  | CDTr | **0.844** | **0.790** | **0.773** | **0.889** | **0.926** | **0.814** |
| 200 | SPIEC(MB) | 0.820 | 0.761 | 0.745 | 0.884 | 0.880 | 0.780 |
|  | SPIEC(GL) | 0.856 | 0.791 | 0.778 | 0.909 | 0.921 | 0.820 |
|  | gCoda | 0.876 | 0.806 | 0.800 | 0.913 | 0.955 | 0.836 |
|  | aCDTr | 0.870 | 0.809 | 0.792 | 0.917 | 0.945 | 0.835 |
|  | CDTr | **0.883** | **0.821** | **0.811** | **0.923** | **0.955** | **0.849** |

### 3.2 Simulations for DCDTr Loss

We investigate the performance of DCDTr loss with some experiments in this section. The first precision matrix Θ is generated as follows:

1. *Random graph*: For Θ, two nodes are connected with probability 0.5, and the strength is generated from a uniform distribution in [−0.4, − 0.2] ∪ [0.2, 0.4].
2. *Band graph*: Connect pair (*i*, *j*) with strength uniformly distributed in [0.05*m* − 0.3, 0.05*m* − 0.25] ∪ [0.25 − 0.05*m*, 0.3 − 0.05*m*], if |*i* − *j*| = *m*, *m* = 1, 2, 3, 4.
3. *Neighbor graph*: Select *p* points from 𝓤(0, 1) and connect the 10 nearest neighbors for each point with strength sampled from a uniform distribution in [−0.4, − 0.2] ∪ [0.2, 0.4].
4. *Scale*-*free graph*: The scale-free graph is generated with the B-A algorithm [4]. The strength between connected nodes is generated from a uniform distribution in [−0.4, − 0.2] ∪ [0.2, 0.4].
5. *Hub graph*: Partition the nodes into 3 disjoint groups evenly and select a node as hub for each group. The hubs are connected with the non-hubs in the same group with strength uniformly distributed in [−0.4, − 0.2] ∪ [0.2, 0.4].
6. *Block graph*: Divide *p* nodes into 5 blocks evenly. Connect pairs in the same block with probability 0.5 and pairs in different blocks with probability 0.3. The strength between connected nodes is uniformly distributed in [−0.4, − 0.2] ∪ [0.4, 0.2].

Then 10% of the connected pairs in Θ will change to an unconnected state, while the same number of unconnected pairs in Θ will change to a connected state, such that we get another precision matrix Θ^∗^. For scale-free and hub graph, the ratio of change is 40% based on the sparsity of the two graphs. The diagonal elements of Θ and Θ^∗^ are set as |*e*| + 0.3, where *e* is the smallest eigenvalue of Θ or Θ^∗^, respectively. The deviations from the exchangeable condition of Θ and Θ^∗^ are listed in Table 5. Therefor, the differential matrix Δ is Θ^∗^ − Θ. The two precision matrices Θ and Θ^∗^ are used to generate data separately. The aforementioned four methods, including DCDTr, FGL, GGL and *ℓ*-M, are used to estimate the true differential matrix Δ. Similarly, we evaluate the true positive rate and true negative rate at different tuning parameters and then compute the area under the ROC curve (AUC). We take the sample size *n* = 100, 200, 300, 400 and repeat this procedure 100 times.

**Table 5.** Deviations from the exchangeable condition of six different network structures.

| Network | Random | Band | Neighbor | Scale-free | Hub | Block |
| --- | --- | --- | --- | --- | --- | --- |
| $dev$ | 0.56 | 0.56 | 0.89 | 0.36 | 1.07 | 1.23 |
| $dev^*$ | 0.48 | 1.02 | 1.03 | 0.39 | 0.49 | 0.99 |

Table 6 presents the mean AUC scores of different methods under different settings. We see that no method is generally better than the others in all cases. DCDTr performs better than other methods in random graph, neighbor graph and block graph, while GGL achieves higher AUC in scale-free and hub graph. With the increase of sample size, the advantage of DCDTr becomes increasingly significant. Generally speaking, our proposed DCDTr performs well in different network estimations.

**Table 6.**
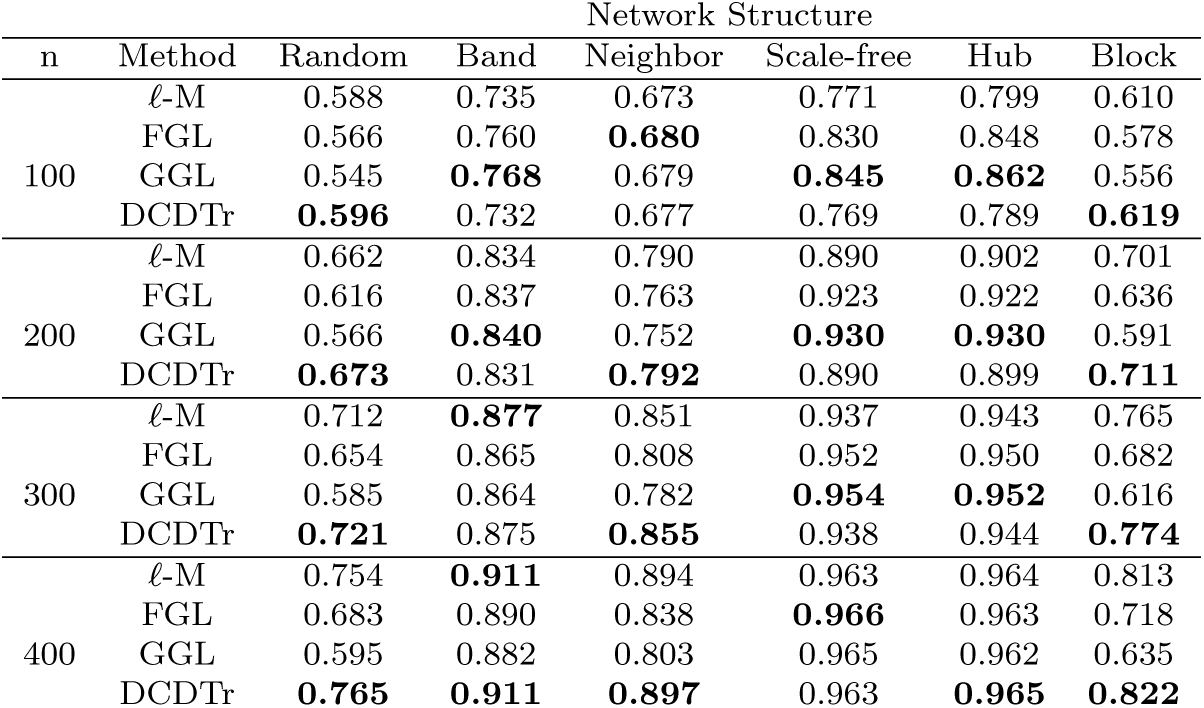
The mean AUC scores of different methods under different settings.

| n | Method | Network Structure |  |  |  |  |  |
| --- | --- | --- | --- | --- | --- | --- | --- |
|  |  | Random | Band | Neighbor | Scale-free | Hub | Block |
| 100 | $\ell$-M | 0.588 | 0.735 | 0.673 | 0.771 | 0.799 | 0.610 |
|  | FGL | 0.566 | 0.760 | **0.680** | 0.830 | 0.848 | 0.578 |
|  | GGL | 0.545 | **0.768** | 0.679 | **0.845** | **0.862** | 0.556 |
|  | DCDTr | **0.596** | 0.732 | 0.677 | 0.769 | 0.789 | **0.619** |
| 200 | $\ell$-M | 0.662 | 0.834 | 0.790 | 0.890 | 0.902 | 0.701 |
|  | FGL | 0.616 | 0.837 | 0.763 | 0.923 | 0.922 | 0.636 |
|  | GGL | 0.566 | **0.840** | 0.752 | **0.930** | **0.930** | 0.591 |
|  | DCDTr | **0.673** | 0.831 | **0.792** | 0.890 | 0.899 | **0.711** |
| 300 | $\ell$-M | 0.712 | **0.877** | 0.851 | 0.937 | 0.943 | 0.765 |
|  | FGL | 0.654 | 0.865 | 0.808 | 0.952 | 0.950 | 0.682 |
|  | GGL | 0.585 | 0.864 | 0.782 | **0.954** | **0.952** | 0.616 |
|  | DCDTr | **0.721** | 0.875 | **0.855** | 0.938 | 0.944 | **0.774** |
| 400 | $\ell$-M | 0.754 | **0.911** | 0.894 | 0.963 | 0.964 | 0.813 |
|  | FGL | 0.683 | 0.890 | 0.838 | **0.966** | 0.963 | 0.718 |
|  | GGL | 0.595 | 0.882 | 0.803 | 0.965 | 0.962 | 0.635 |
|  | DCDTr | **0.765** | **0.911** | **0.897** | 0.963 | **0.965** | **0.822** |

## 4 Real Data Analysis

In this section, we illustrate our proposed method with an application to mouse skin microbiome data [23]. A total of 261 mice were divided into 3 groups: 78 non-immunized controls (Control), 119 immunized healthy individuals (Healthy) and 64 immunized epidermolysis bullosa acquisita individuals (EBA), according to the health conditions of skin immunizations. The OTUs appearing in less than 50% of the samples are filtered out, and the samples with a number of nonzero OTU counts less than 50% of the total selected OTUs are also removed. We finally arrived at a dataset with *p* = 77 OTUs and *n* = 232 samples (63 Control, 114 Healthy and 55 EBA). We increased all OTU counts by 0.5 to avoid zero counts and normalized the data to compositional data.

Since the the underlying true direct interaction networks were not available and the accuracy of estimated networks was unobtainable, we evaluated the performance of the proposed methods with reproducibility as suggested by Fang *et al.* [10] and Kurtz *et al.* [18] suggusted. More specifically, we first constructed a reference network *est*_1_ (precision matrix or differential matrix) with all data for each group and method. We then selected half of the samples randomly to estimate the precision matrix or differential matrix (denoted by *est*_2_) again. The reproducibility was measured by the fraction of overlapping edges shared by *est*_1_ and *est*_2_ in the reference network *est*_1_.

For each group and each method of precision matrix estimation, the procedure stated above was repeated 20 times. The mean reproducibility is summarized in Table 7. CDTr and aCDTr outperformed the other three methods in terms of reproducibility in all three groups, implying that CDTr and aCDTr are more stable and accurate in direct interaction estimation. We also estimated the differential network for the Control-Healthy and Control-EBA groups, and the evaluation procedure was also repeated 20 times. The mean reproducibility is listed in Table 8. The highest reproducibility of DCDTr also implies that DCDTr is more stable and accurate in differential network estimation.

**Table 7.** The mean reproducibility for various methods and groups.

|  | SPIEC(MB) | SPIEC(GL) | gCoda | aCDTr | CDTr |
| --- | --- | --- | --- | --- | --- |
| Control | 0.47 | 0.58 | 0.29 | 0.60 | 0.65 |
| Healthy | 0.57 | 0.70 | 0.15 | 0.84 | 0.84 |
| EBA | 0.51 | 0.57 | 0.11 | 0.94 | 0.93 |

**Table 8.** The mean reproducibility for various methods and groups.

| | $\ell_1$-M | FGL | GGL | DCDTr |
| --- | --- | --- | --- | --- |
| Control-Healthy | 0.45 | 0.54 | 0.56 | 0.78 |
| Control-EBA | 0.50 | 0.65 | 0.67 | 0.84 |

Finally, we employed all methods to build a candidate microbiome association network from the unified dataset for each group and group pairs. In Fig **??**, we present the number of shared edges for direct interaction networks recovered from various methods via Venn diagrams. We can see that the direct interaction network from CDTr is close to that of aCDTr, while the network from SPIEC(GL) and SPIEC(MB) are more similar. A total of 28, 46 and 24 edges are shared by all candidate networks for control, healthy and EBA groups, respectively, comprising the core interaction network among OTUs. Moreover, almost all direct interactions discovered by CDTr and aCDTr are in this core interaction network, while SPIEC(GL), SPIEC(MB) and gCoda discover some eccentric interactions. The number of shared edges for differential networks are shown in Fig **??**. The situation for differential networks is much more complicated. *ℓ*_1_-M discovered many eccentric differential edges in both groups, but these were not confirmed by other methods. The differential edges from GGL and FGL are almost the same for both groups, and are more than the edges from DCDTr. Most differential edges from DCDTr were verified by both GGL and FGL for both groups, implying that DCDTr is good at inferring the crucial differential edges without mixing nonessential edges.

## 5 Conclusion

Inferring the direct interactions among microbial species and understanding how the network structure changes are important in the study of ecology and medicine. In this paper, we propose a framework to estimate the direct interaction network and differential network from compositional microbial data based on clr transformation and D-trace loss for absolute abundance data. Although the proposed CDTr loss and DCDTr loss are derived from an exchangeable condition, we show that they still perform well and better than other methods under different scenarios in our numerical simulations. However, the reasonableness of the exchangeable condition should be further examined in theory and biology. Finally, the consistency of the estimators does not come with a theoretical guarantee, which is a common limitation of gCoda, SPIEC, CDTr and DCDTr. For future work, we are interested in developing theorems about the consistency property in both direct interaction network and differential network estimation.

## Supporting information

## S1 Appendix. Supplementary for A Framework to Incorporate D-trace Loss into Compositional Data Analysis.

The matrix operators *S*(*X*),*K*(*X*),*H*(*X*) and [*X*]_+_ used in Algorithm 1 and Algorithm 2 for the numerical solutions of lasso penalized CDTr and DCDTr loss are presented in this Supplementary. We also demonstrate the relationship between D-trace loss and CDTr loss, as well as the relationship between DTL loss and DCDTr loss. The detailed formulas of *ℓ*_1_-minimization method and joint graphical lasso (FGL, GGL) are listed in this Supplementary.

## Acknowledgments

This work was supported by The National Key Research and Development Program of China (No.2016YFA0502303), the National Key Basic Research Project of China (No. 2015CB910303), and the National Natural Science Foundation of China (No.31471246).

---

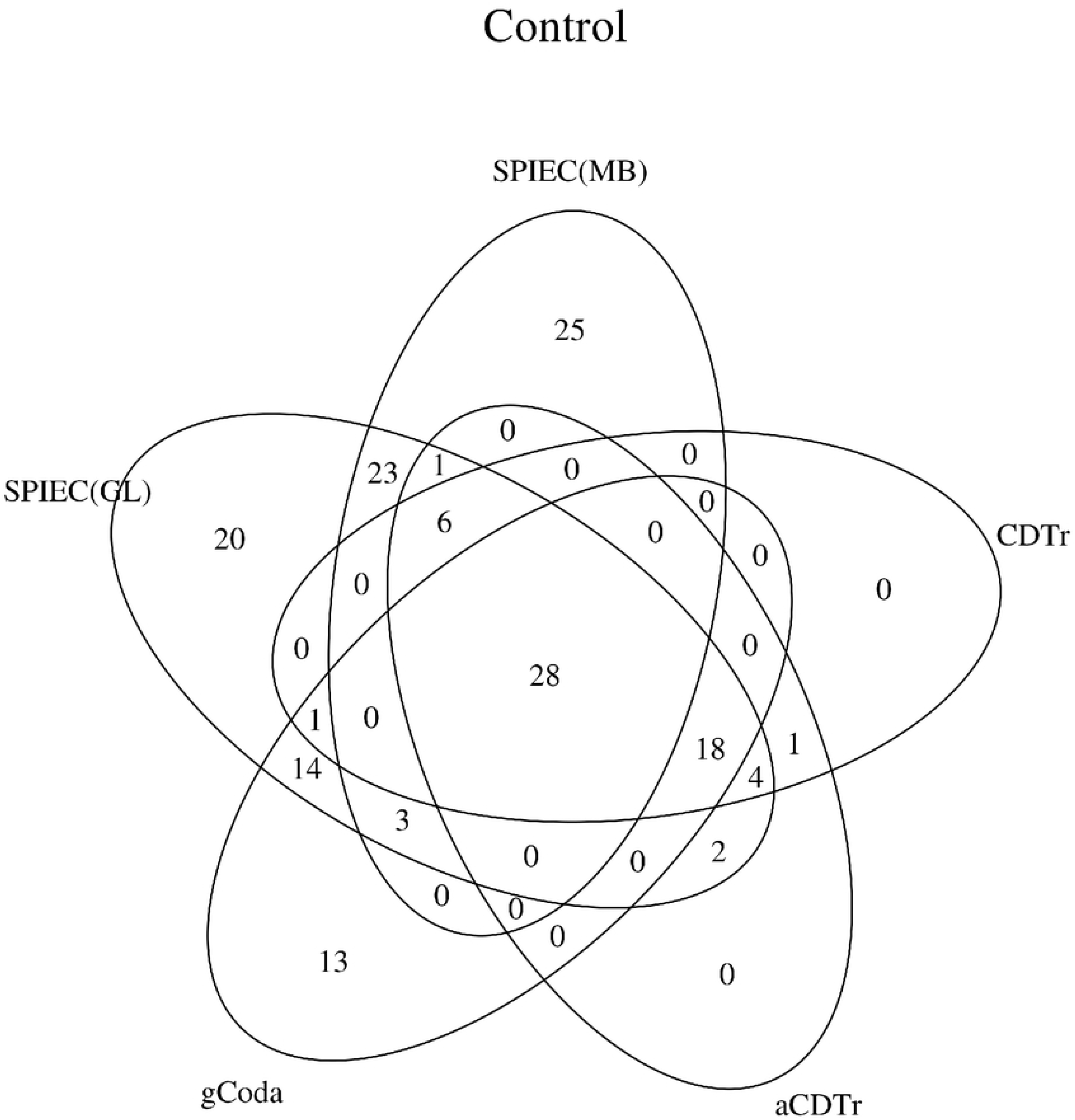

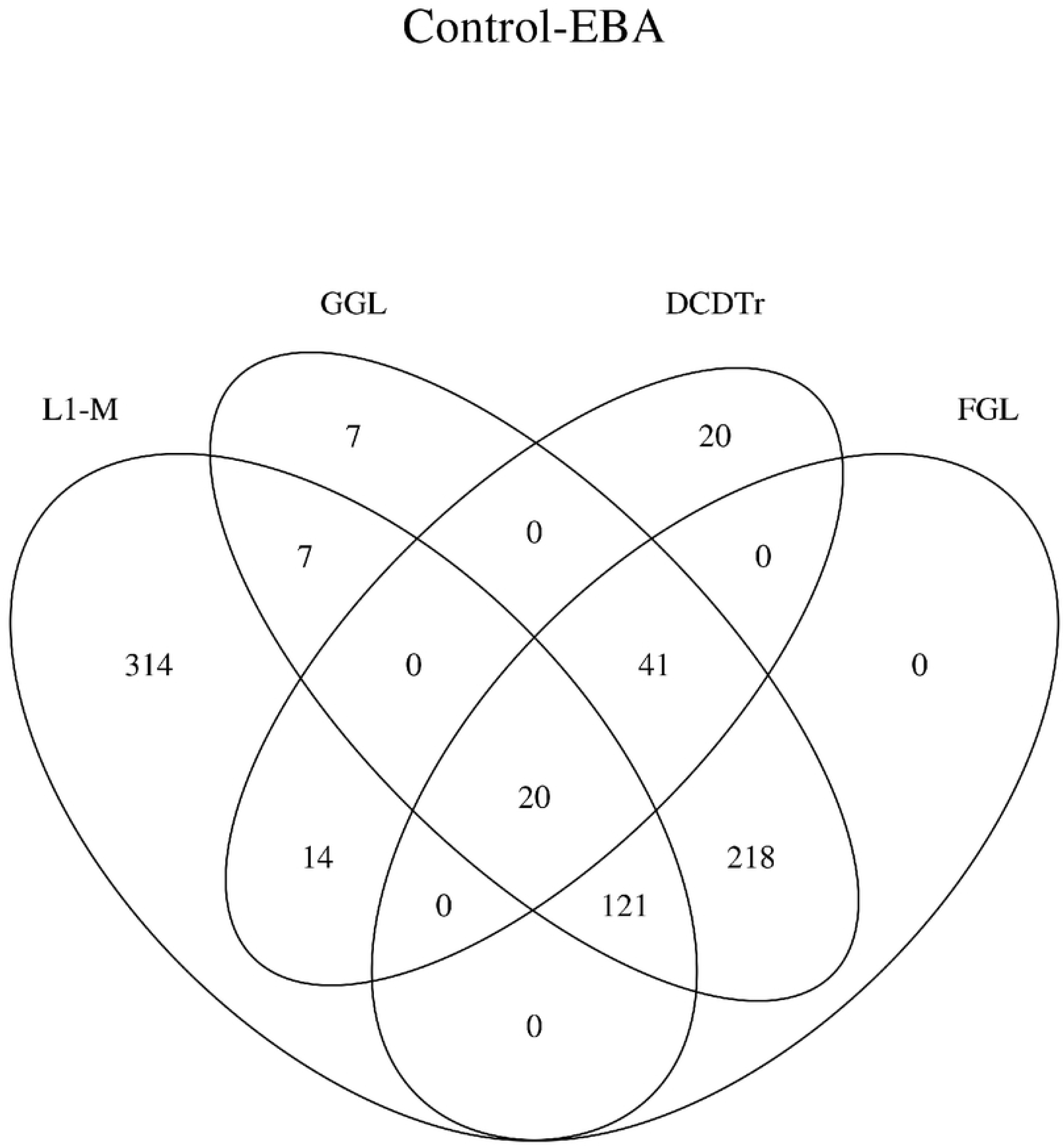

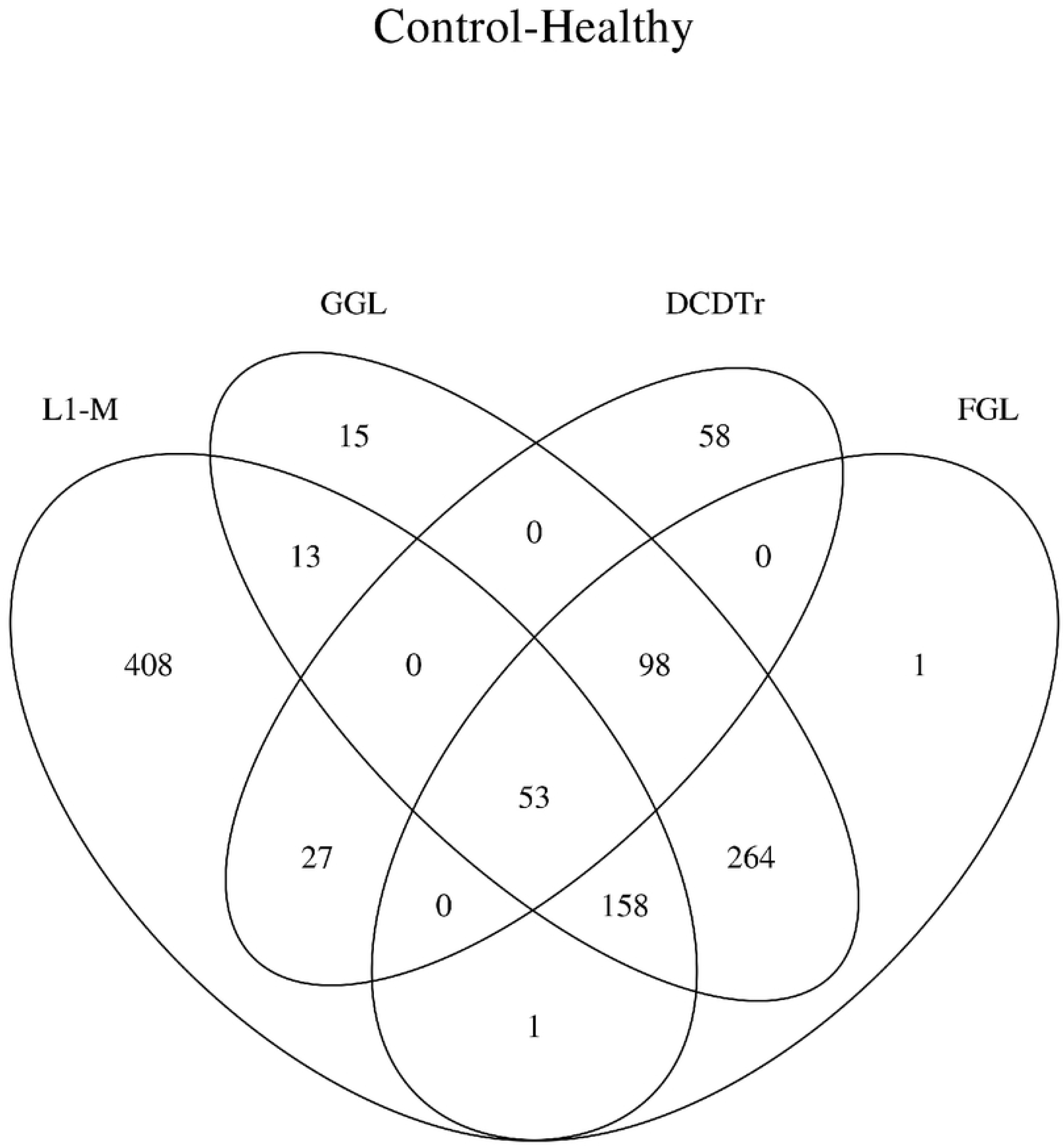

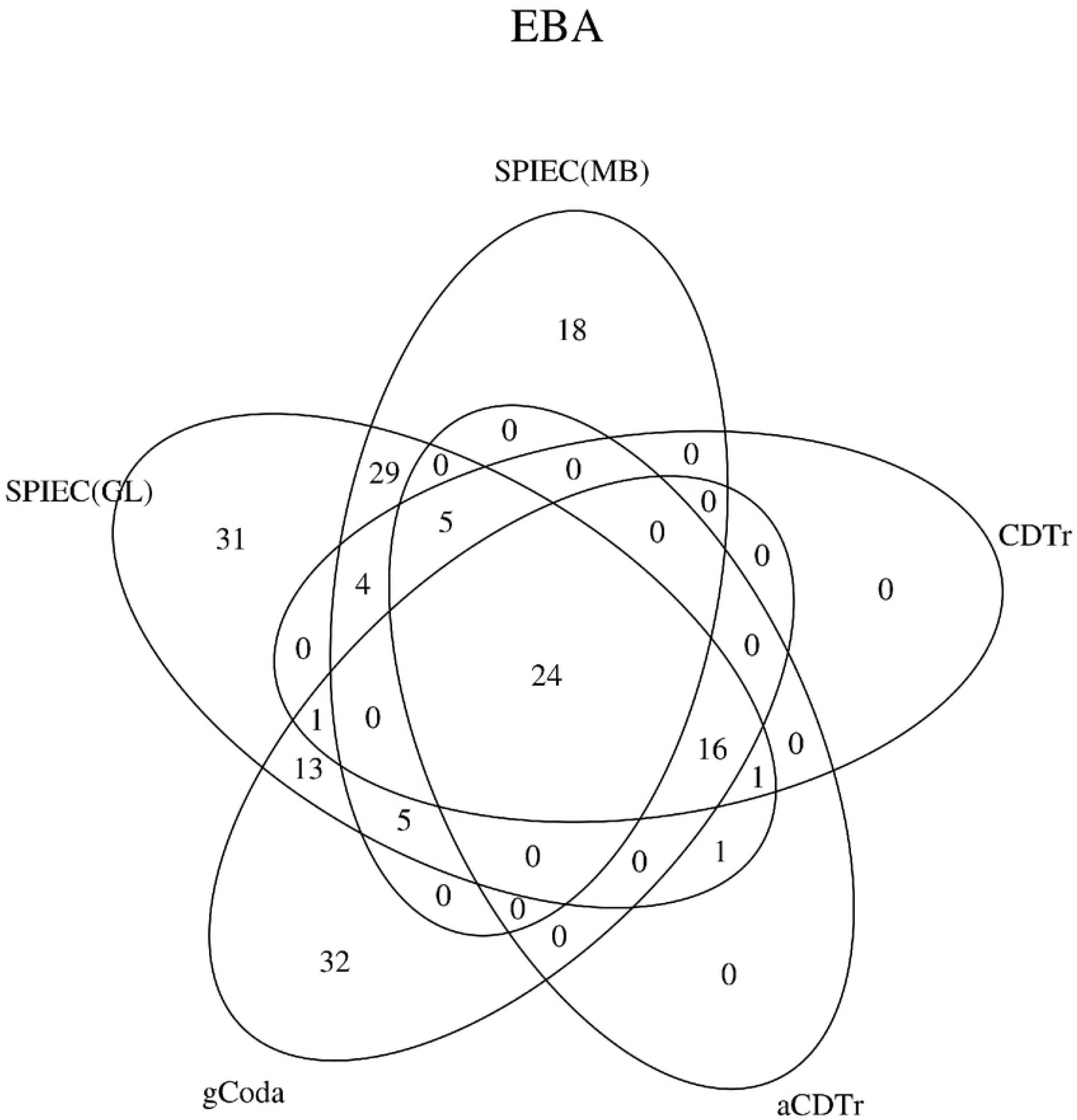

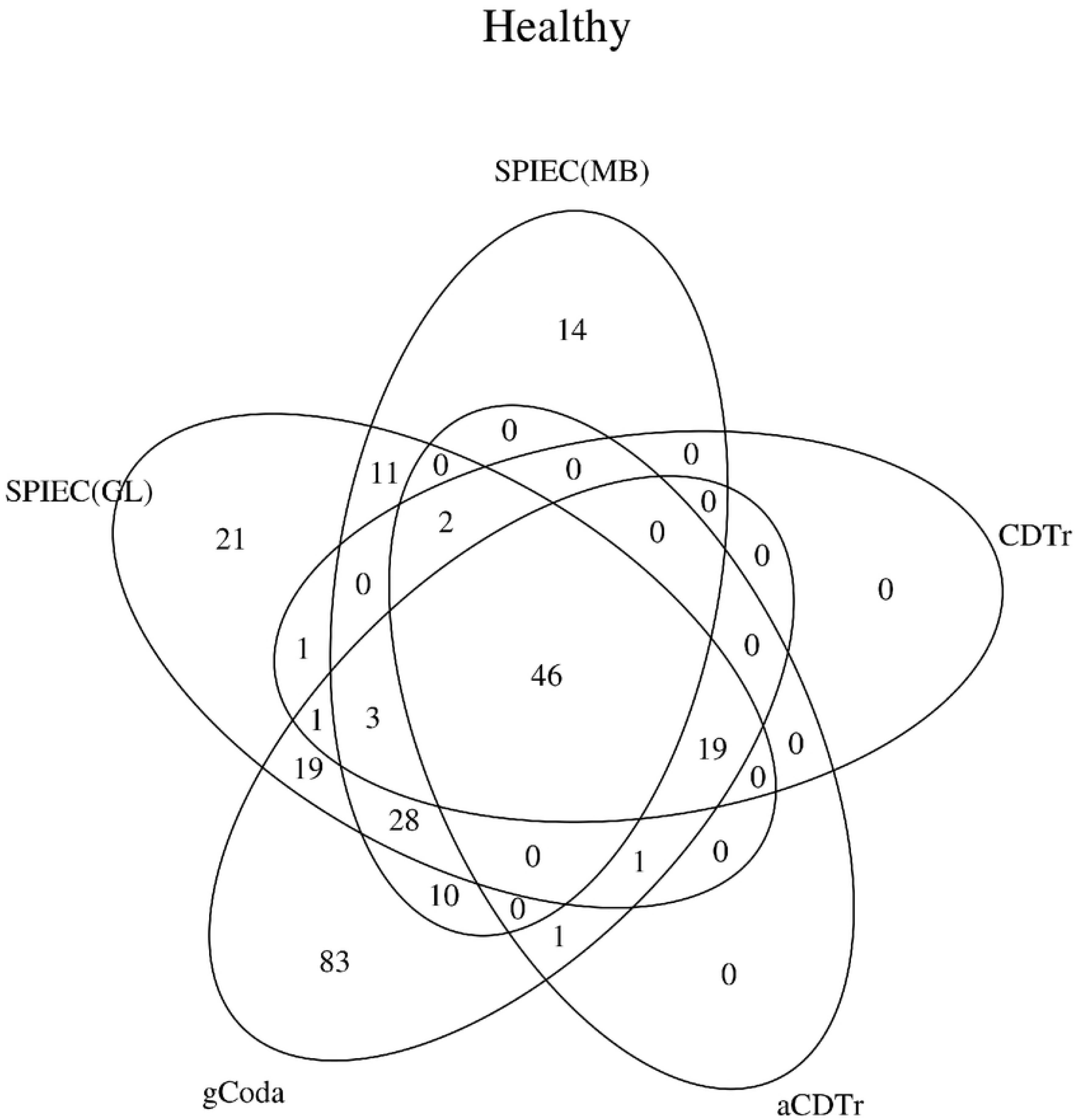

## References

1. John Aitchison. A new approach to null correlations of proportions. Mathematical Geology, 13(2):175–189, 1981.

2. Yuguang Ban, Lingling An, and Hongmei Jiang. Investigating microbial co-occurrence patterns based on metagenomic compositional data. Bioinformatics, 31(20):3322–3329, 2015.

3. Sourav Bandyopadhyay, Monika Mehta, Dwight Kuo, Min-Kyung Sung, Ryan Chuang, Eric J Jaehnig, Bernd Bodenmiller, Katherine Licon, Wilbert Copeland, Michael Shales, et al. Rewiring of genetic networks in response to dna damage. Science, 330(6009):1385–1389, 2010.

4. Albert-László Barabási and Reka Albert. Emergence of scaling in random networks. science, 286(5439):509–512, 1999.

5. Surojit Biswas, Meredith Mcdonald, Derek S. Lundberg, Jeffery L. Dangl, and Vladimir Jojic. Learning microbial interaction networks from metagenomic count data. Journal of Computational Biology A Journal of Computational Molecular Cell Biology, 23(6):526, 2016.

6. Julien Chiquet, Yves Grandvalet, and Christophe Ambroise. Inferring multiple graphical structures. Statistics and Computing, 21(4):537–553, 2011.

7. Patrick Danaher, Pei Wang, and Daniela M. Witten. The joint graphical lasso for inverse covariance estimation across multiple classes. Journal of the Royal Statistical Society: Series B (Statistical Methodology), 76(2):373–397, 2014.

8. P. G. Falkowski, T Fenchel, and E. F. Delong. The microbial engines that drive earth’s biogeochemical cycles. Science, 320(5879):1034–9, 2008.

9. Huaying Fang, Chengcheng Huang, Hongyu Zhao, and Minghua Deng. Cclasso: correlation inference for compositional data through lasso. Bioinformatics, 31(19): 3172–3180, 2015.

10. Huaying Fang, Chengcheng Huang, Hongyu Zhao, and Minghua Deng. gcoda: Conditional dependence network inference for compositional data. Journal of Computational Biology, 2017.

11. Karoline Faust and Jeroen Raes. Microbial interactions: from networks to models. Nature reviews. Microbiology, 10(8):538, 2012.

12. Karoline Faust, J Fah Sathirapongsasuti, Jacques Izard, Nicola Segata, Dirk Gevers, Jeroen Raes, and Curtis Huttenhower. Microbial co-occurrence relationships in the human microbiome. PLoS computational biology, 8(7):e1002606, 2012.

13. Jerome Friedman, Trevor Hastie, and Robert Tibshirani. Sparse inverse covariance estimation with the graphical lasso. Biostatistics, 9(3):432–441, 2008.

14. Jonathan Friedman and Eric J Alm. Inferring correlation networks from genomic survey data. PLoS computational biology, 8(9):e1002687, 2012.

15. Nir Friedman. Inferring cellular networks using probabilistic graphical models. Science, 303(5659):799–805, 2004.

16. Jian Guo, Elizaveta Levina, George Michailidis, and Ji Zhu. Joint estimation of multiple graphical models. Biometrika, 98(1):1–15, 2011.

17. Allan Konopka. What is microbial community ecology—[quest]—. Isme Journal, 3(11):1223, 2009.

18. Zachary D Kurtz, Christian L Müller, Emily R Miraldi, Dan R Littman, Martin J Blaser, and Richard A Bonneau. Sparse and compositionally robust inference of microbial ecological networks. PLoS computational biology, 11(5):e1004226, 2015.

19. Florian Markowetz and Rainer Spang. Inferring cellular networks–a review. BMC bioinformatics, 8(6):S5, 2007.

20. Nicolai Meinshausen and Peter Bühlmann. High-dimensional graphs and variable selection with the lasso. The annals of statistics, pages 1436–1462, 2006.

21. Katya Scheinberg, Shiqian Ma, and Donald Goldfarb. Sparse inverse covariance selection via alternating linearization methods. Advances in Neural Information Processing Systems, pages 2101–2109, 2010.

22. Gideon Schwarz et al. Estimating the dimension of a model. The annals of statistics, 6(2):461–464, 1978.

23. Girish Srinivas, Steffen Möller, Jun Wang, Sven Künzel, Detlef Zillikens, John F Baines, and Saleh M Ibrahim. Genome-wide mapping of gene–microbiota interactions in susceptibility to autoimmune skin blistering. Nature communications, 4, 2013.

24. I Thiele, A Heinken, and R. M. Fleming. A systems biology approach to studying the role of microbes in human health. Current Opinion in Biotechnology, 24(1): 4–12, 2013.

25. Robert Tibshirani. Regression shrinkage and selection via the lasso. Journal of the Royal Statistical Society. Series B (Methodological), pages 267–288, 1996.

26. Joe Whittaker. Graphical models in applied multivariate statistics. Wiley Publishing, 2009.

27. John C Wooley, Adam Godzik, and Iddo Friedberg. A primer on metagenomics. PLoS computational biology, 6(2):e1000667, 2010.

28. Huili Yuan, Ruibin Xi, and Minghua Deng. Differential network analysis via the lasso penalized d-trace loss. Biometrika, 104(4), 2015.

29. Ming Yuan and Yi Lin. Model selection and estimation in the gaussian graphical model. Biometrika, 94(1):19–35, 2007.

30. Teng Zhang and Hui Zou. Sparse precision matrix estimation via lasso penalized d-trace loss. Biometrika, 101(1):103–120, 2014.

31. Sihai Dave Zhao, T. Tony Cai, and Hongzhe Li. Direct estimation of differential networks. Biometrika, 101(2):253–268, 2014.

